# An in-silico comparative analysis of lncRNA expression and their role in the pathogenesis of representative fungal, bacterial and viral infections in rice

**DOI:** 10.1101/2024.08.18.608517

**Authors:** Manu Kandpal, Mahi Sharma, Bhadresh R. Rami

## Abstract

Long non-coding RNAs (lncRNAs) perform prominent role in the regulation of gene expression during plant development and stress response by directly interacting with DNA, RNA, proteins, and/or triggering production of small regulatory RNA molecules. The objective of our study is to understand the systems-level response of the same plant species to highly diverse pathogens across different kingdoms and evaluate the patterns of similarity vs differences, specifically in the context of lncRNA’s. Towards this objective, we performed a comparative in silico analysis of lncRNA’s of Rice that are differentially expressed in response to infection by bacteria (*Xanthomonas oryzae*), fungus (*Magnaporthe oryzae*) and virus (*Rice black dwarf virus*). Using a tailored lncRNA analysis pipeline, we successfully identified 1125, 719 and 240 lncRNAs in *Xanthomonas oryzae* infection susceptible cultivar CT9737-6-1-3P-M, *Magnaporthe oryzae* susceptible LTH accession, and *Rice black streaked dwarf virus* susceptible Wuyujing No. 7 rice cultivars respectively. The in-silico predicted Cis- and Trans-target genes of lncRNAs were subsequently used to identify the pathways modulated by these lncRNA and how they cluster into unique categories of plant responses to pathogen infections. To further substantiate the role of predicted lncRNA’s in plant defence and immune response our analysis finds that many of the lncRNAs co-localize with the QTLs associated with Blast and Bacterial blight resistance in rice. Our in silico analysis provides a list of common and unique pathogen specific lncRNAs that can provide vital insights into the generic vs tailored mechanisms adopted by rice in different infection scenarios.

## 1 Introduction

Ensuring a stable global food supply is vital for human well-being and the sustainability of civilization, especially with the increasing impact of climate change. Rice, wheat, maize, and potatoes are key staples in the global diet. Rice makes up approximately 8% (FAO. 2022) of global crop production. Rice is the primary staple in Asia, the Caribbean, and Latin America. Potatoes are central to European diets (Oecd 2022) making up 4.14% of global crop production (FAO 2022). Maize contributes to 12% (FAO. 2022) of global crop production and dominates in the Americas, where it constitutes 50% (FAOStat. 2021) of maize production. In Africa, especially in the east and south, maize is crucial for food and nutrition security (FAOStat. 2021). Likewise, wheat, contributing 8% of the world’s total crop production, stands as a dietary staple in Northern Africa, West/Central Asia, and Europe (Erenstein et al. 2022).

Rice is a vital staple crop, essential for global food security and livelihoods. It is extensively grown in countries like China, India, Indonesia, Bangladesh, Vietnam, and Thailand (FAOStat. 2021). Rice is a primary source of energy and nutrients, particularly in Asia, deeply rooted in cultural traditions and diets. Its cultivation offers employment, income, and economic contributions. Ensuring access to diverse and affordable rice varieties is crucial for fighting hunger and establishing sustainable food systems, making rice not just a food crop but also a symbol of tradition and prosperity.

Loss in the yield of rice crop due to pathogens and pests can range from ∼25%-41% (Savary et al. 2019), posing a significant threat to food security. Global warming further exacerbates the problem by weakening plant immunity and altering pathogen prevalence and infection patterns. Elevated temperatures make plants more susceptible to pathogens, leading to higher disease incidence. Prioritizing crop health management and adopting integrated approaches are essential for sustainable agriculture. Understanding pathogen infection mechanisms, including plant responses and pathogen strategies, is crucial. In this study, we investigate three pathogens affecting rice: Bacteria, *Xanthomonas oryzae* (causing 10-50% yield loss) (Lee et al. 2005; Liu and Wang 2016), Fungus, *Magnaporthe oryzae* (causing up to 100% yield loss) (Dean et al. 2005; Liu and Wang 2016), and a virus, *Rice black streaked dwarf virus* (causing ∼60% yield loss) (Liu and Wang 2016). Studying these pathogens helps us grasp the dynamics of plant-pathogen interactions across highly diverse pathogens and the strategies employed by host plant *vs* the pathogens.

Long non-coding RNAs (lncRNAs) play crucial roles in cellular regulation, revolutionizing our understanding of gene control. These long RNA molecules, with a typical length >200 nucleotides, lack protein-coding capacity are involved in diverse biological processes (Rinn and Chang 2012). They function in scaffolding (Yoon et al. 2013), epigenetic regulation (Jain et al. 2016), transcriptional regulation (Feng et al. 2006; Li et al. 2014), post-transcriptional regulation (Bardou et al. 2014), miRNA Sponging (Thomson and Dinger 2016), and cellular signalling & subnuclear compartmentalization (Clemson et al. 2009).

In plants, lncRNAs are significant regulators influencing flowering time (Berry and Dean 2015), management of abiotic (Zhang et al. 2019) and biotic (Zaynab et al. 2021) stresses, development of leaf (X et al. 2018), root growth (Rigo et al. 2020) and reproduction (Wu et al. 2013).

Our in silico study aims to systematically compare the effects of three kingdoms of pathogens, i.e., bacteria, fungi, and viruses on a single crop, rice, using available transcriptomics data. We aim to understand the role of lncRNA regulation during pathogenic infections by the bacteria, *Xanthomonas oryzae* (henceforth referred to Xoo), fungus, *Magnaporthe oryzae* (henceforth referred to Mor), and virus, *Rice black streaked dwarf virus* (henceforth referred to RBSDV). Our study reveals molecular insights into how common and unique lncRNAs influence response of rice to various agricultural pathogens thus opening new avenues for understanding complex plant-pathogen interactions.

## 2 Methods

### 2.1 Data collection and data analysis

RNA sequencing data were obtained from NCBI GEO PRJNA631554 (Shu et al. 2021) for R-B (Rice species CT-9737-6-1-3P-M + Bacteria (Xoo)), PRJNA545418 (Fan et al. 2020) for (Rice species LTH + Fungus (Mor)), and PRJNA657713 (Zhang et al. 2020a)) for (Rice species Wuyujing No.7 + Virus (RBSDV)) from the leaf samples of susceptible rice species. Table 1 shows the three pairs of infections considered in this study.

**Table 1.** List of pathogens and rice cultivars used in this study, along with site and duration of infection and impact on yield.

| # | Pathogen | Type | Rice cultivars | Tissue | Duration of infection | Impact (↓ yield) |
| --- | --- | --- | --- | --- | --- | --- |
| 1 | Xanthomonas oryzae pv. oryzae | Bacteria | CT 9737-6-1-3P-M | Leaf | 24 hrs post infection | 10-50% |
| 2 | Magnaporthe oryzae | Fungi | LTH (blast susceptible japonica accession) | Leaf | 24 hrs post infection | Up to 100% |
| 3 | Rice black streaked dwarf virus | Virus | Wuyujing No. 7 | Leaf | Infection was given at three leaf stage till the onset of disease | ~60% |

RNA-seq raw data were sourced from previous studies for all three R-B, R-F, and R-V infections. In these studies, RNA sequencing was performed using three biological replicates of each infected leaf 24 h after infection with fungi and bacteria. RNA sequencing data for virus-infected leaves were taken 1-week after infection. In the case of R-F & R-B, only transcriptomics analysis was performed, without any focus on lncRNA. In the R-V combination, while identification of lncRNA was carried out, however, the tools used for the analysis were different. Zhang et al. (2020a) used Tophat2 for alignment and Cuffdiff for differential expression, while in our analysis, we used Hisat2 and DESeq2. Hisat2 is better than Tophat2 in speed, efficiency, and accuracy. Similarly, DESeq2 offers advantages over Cuffdiff, such as improved accuracy in estimating gene expression variability, better handling of low count genes, and a robust framework for detecting differential expression with reduced false positive rates.

Therefore we subjected all the three data sets to consistent and uniform analysis methods to ensure reliable and reproducible results by minimizing technical variability and standardizing the workflow and focused specifically towards lncRNA identification and characterization.

Raw samples in fastq format were filtered to remove adapter-contaminated and low-quality reads. Hisat2 (Kim et al. 2015) was then used to align these high-quality reads with a Phred score ≥30 to the rice genome obtained from the Rice Genome Annotation Project Database version 7. Assembly was performed using a stringtie (Pertea et al. 2015). The final merged assembly was prepared using the Cuffmerge (Trapnell et al. 2012) software. The read count for all the identified transcripts was calculated using bedtools.

### 2.2 Identification and expression analysis of lncRNA’s

We utilized a computational analysis pipeline, as shown in Figure 1, to identify lncRNAs in rice species infected by fungi, bacteria, and viruses. Sequences were extracted from merged assemblies using GffRead (Pertea and Pertea 2020) to isolate lncRNAs from the total transcripts. The longest transcript for each locus was then subjected to length filtering using a Perl script. These transcripts were further filtered based on the class code (i-intronic, u-intergenic, and x-antisense). Transcripts with >200 nt length were selected and shorter transcripts were removed.

**Figure 1.**
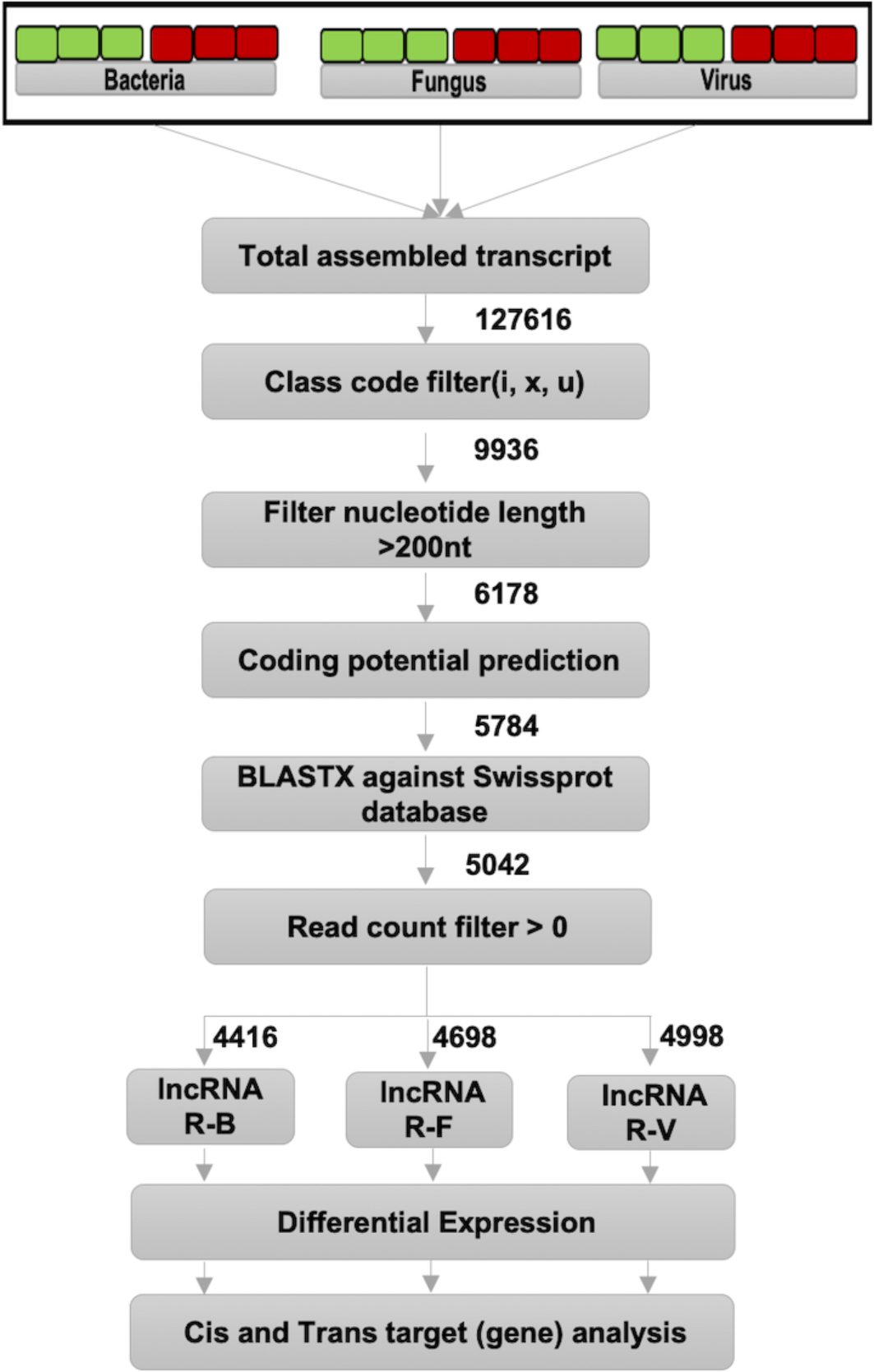
Workflow for lncRNA identification: Flow chart depicts the steps for identification lncRNA. Numbers next to arrows reflect the filtration happening at each step of the analysis pipeline based on the cutoff criteria employed with respect to lncRNA identification. Transcripts of LTH, CT 9737-6-1-3P-M, and Wuyujing #7 cultivars of rice which were infected with Magnaporthe oryzae, Xanthomonas oryzae pv. Oryzae, and Rice black streaked dwarf virus, respectively, the control and infected samples were obtained from different studies mentioned in section 2.1. R-B: Xanthomonas oryzae + CT9737-6-1-3P-M, R-F: Magnaporthe oryzae + LTH and R-V: Rice black streaked dwarf virus + Wuyujing No. 7

The coding potential of all remaining transcripts was assessed using the coding potential calculator (CPC2) (Kang et al. 2017) with a threshold score of <0.5. These sequences were subsequently filtered based on alignment with the SwissProt Database (Magrane and Consortium 2011) using BLASTX (E-value < 0.001) and checked for the occurrence of Pfam protein domains (Finn et al. 2014) with Hmmscan (E-value < 0.001).

Following the completion of the filtration process, transcripts that successfully meet all criteria are designated as lncRNAs. Subsequently, the expression of these lncRNAs were confirmed based on their read counts being >0. The differential expression between control and infected samples was determined using Deseq2 (Love et al. 2014). The cis and trans genes analysis in Figure 1 is elaborated in section 2.4.

### 2.3 Mapping lncRNAs overlapping Quantitative trait loci (QTL)

We investigated whether DE-lncRNAs in plants infected by bacteria, fungi, and viruses overlap with previously identified QTLs listed in RiceLncPedia (Zhang et al. 2020b). To visualize the spatial distribution of QTLs and lncRNAs on rice chromosomes, we utilized ChromoMap (Figure 3) (Anand and Rodriguez Lopez 2022).

### 2.4 Target gene prediction and functional analysis of the target DE-genes

LncRNAs can regulate protein-coding genes either locally (cis-regulation) or at a distance (trans-regulation). In our study we identified protein-coding genes located within 100 kb of lncRNAs as potential cis-regulation targets. For trans-regulation, we calculated the Pearson correlation coefficient (r) between differentially expressed lncRNAs (DE-lncRNAs) and differentially expressed genes (DE-genes). An r value of ≥0.80 and a significance level (P-value) ≤0.00001 indicate potentially strong trans-regulation relationships between lncRNAs and genes (Figure 5A). These cis/trans effects of DE-lncRNAs can lead to gene upregulation or downregulation, depending on their specific mechanisms.

The expression values (calculated in terms of transcript per million (TPM)) for all the lncRNA’s and genes were used to perform Principal component analysis (PCA) using prcomp function of the R package stats (Figure 4).

KOBAS-i (Bu et al. 2021) was used to perform KEGG pathway enrichment analysis to identify which specific biological pathways are influenced by the lncRNAs. Pathways with a P-value ≤0.05 were considered as significantly enriched (Figure 5B).

## 3 Results

### 3.1 Identification and characterization of lncRNA

In Figure 1, we illustrate our computational analysis workflow for identifying lncRNAs in infected and control transcriptome data (R-B, R-F, and R-V infections). Initially, 127,616 aligned transcripts were used to identify lncRNAs. Using a class code-based approach, 9,936 transcripts were retained after filtering out coding transcripts. The class code system distinguishes coding from non-coding transcripts based on genomic location: "x" for exonic overlap, "=" for complete intron chain match, "i" for intron-contained transfrags, and "u" for intergenic transcripts. After eliminating transcripts <200 nucleotides, 6,178 transcripts remained. Subsequently, CPC2.0 was employed to remove transcripts with protein coding potential, resulting in 5,784 transcripts. These were further analyzed for sequence similarity to coding proteins and the presence of Pfam domains. In our analysis pipeline, we identified 5,042 unique transcripts as lncRNAs, with 3,884 being intergenic, 1,156 antisense, and two intronic. This highlights that the majority of lncRNAs in our study are long *intergenic* non-coding RNAs.

### 3.2 Differential expression analysis of rice lncRNAs upon infection by different pathogens

Table 2 shows that of the 5,042 unique lncRNA’s identified, our analysis identified 4,416 annotated lncRNAs in R-B, 4,698 in R-F, and 4,998 in R-V. A False Discovery Rate (FDR) of ≤0.05, was used to identify whether differentially expressed lncRNA’s are significant or not. In comparison to the uninfected controls, 1124, 691, and 124 lncRNA’s were differentially expressed to a significant extent in R-B, R-F and R-V. Of the 1124 DE-lncRNA’s in R-B, 589 were upregulated and 535 were downregulated. Of the 691 DE-lncRNA’s identified in R-F, 442 and 249 lncRNAs were upregulated and downregulated, respectively. Similarly, of the 240 DE-lncRNA’s in R-V, the upregulated and downregulated lncRNAs were 63 and 177, respectively.

**Table 2:** Overview of total number of lncRNA vs number of differentially expressed (DE) lncRNA’s in the context of different infections.

| # | Pathogen – Plant pair | Total # of lncRNA identified | # of DE lncRNA's | # of upregulated lncRNA | # of downregulated lncRNA |
| --- | --- | --- | --- | --- | --- |
| 1 | Xanthomonas oryzae + CT9737-6-1-3P-M | 4416 | 1124 | 589 | 535 |
| 2 | Magnaporthe oryzae + LTH | 4698 | 691 | 442 | 249 |
| 3 | Rice black streaked dwarf virus + Wuyujing No. 7 | 4993 | 240 | 63 | 177 |

**Table 3.** List of DE-lncRNAs shortlisted based on the functions of target genes implicated in plant pathogen interaction.

| # | DE-lncRNA | Target gene name | Target gene ID | Traits affected | References |
| --- | --- | --- | --- | --- | --- |
| **Common R-B, R-F & R-V** |  |  |  |  |  |
| 1 | TCON_00032885 | OsCHI7 | LOC_Os12g02370 | Regulate flavanoid biosynthesis | Blount et al. 1992;<br>Dai et al. 1996;<br>Beckman 2000 |
|  | TCON_00032893 |  |  |  |  |
|  | TCON_00032892 |  |  |  |  |
| 2 | TCON_00040564 | OsPIL11 | LOC_Os12g41650 | Important for crop yield and plant resistance | Wang et al. 2018 |
| 3 | TCONS_00009600 | OsALMT4 | LOC_Os01g12210 | malate-permeable ion channel with their role in plant pathogen interactions | Liepman and Olsen 2001; Luo et al. 2021 |
| **Common R-B & R-F** |  |  |  |  |  |
| 4 | TCONS_00032659 | OsSWEET14 | LOC_Os11g31190 | sugar transporter, plays a pivotal role in regulating plant susceptibility to pathogens by modulating sugar content | Luo et al. 2021 |
| 5 | TCONS_00072332 | OsbHLH65 | LOC_Os04g41570 | Transcription factor that is phosphorylated by OsMPK3 to induce different defence signal transduction pathways | Shin et al. 2014 |
| 6 | TCONS_00084044 | OsAMTR1 | LOC_Os05g39770 | Aminotransferase enzyme family, plays a crucial role in regulating stress responses during biotic stress | Kothari et al. 2016 |
| 7 | TCONS_00023442 | OsSRO1a | LOC_Os10g42710 | Regulation of JA signaling | Kashihara et al. 2020 |
| 8 | TCONS_00085534 | Xa5 | LOC_Os05g01710 | TAL-mediated immunity | Chen and Ronald 2011 |
| **Common R-B & R-V** |  |  |  |  |  |
| 9 | TCONS_00080140 | OsRR6 | LOC_Os04g57720 | Negative regulator of CK signalling | Hirose et al. 2007 |
| **Common R-F & R-V** |  |  |  |  |  |
| 10 | TCONS_00084595 | OsrbohD | LOC_Os05g45210 | Plays a crucial role in recognizing PAMPs | Kadota et al. 2015 |
| 11 | TCONS_00009525 | OsNCX1 | LOC_Os01g11414 | Sodium/Calcium exchanger (NCX) gene family in rice, regulating calcium ion (Ca2+) | Singh et al. 2015 |
| Unique to R-B |  |  |  |  |  |
| 12 | TCONS_00016268 | OsOSM1 | LOC_Os12g38170 | Resistance against sheath blight, in response to JA | Xue et al. 2016 |
| 13 | TCONS_00097445 | CRK6 | LOC_Os07g35690 | Trigger NH1-mediated immunity | Chern et al. 2016 |
| 14 | TCONS_00090193 | OsJAR1 | LOC_Os05g50890 | Stress-induced JA-Ile | SHIMIZU et al. 2013 |
|  | TCONS_00085279 |  |  |  |  |
|  | TCONS_00085294 |  |  |  |  |
| Unique to R-F |  |  |  |  |  |
| 15 | TCONS_00040972 | OsTrxm | LOC_Os12g08730 | Regulation of ROS levels | Yong et al. 2008 |
|  | TCONS_00040973 |  |  |  |  |
| 16 | TCONS_00054613 | OsPrx30 | LOC_Os02g14430 | Scavenger of ROS | Liu et al. 2021 |
|  | TCONS_00054614 |  |  |  |  |
| 17 | TCONS_00017481 | OsGST4 | LOC_Os01g25100 | Scavenger of ROS | Xu et al. 2018 |
|  | TCONS_00016176 |  |  |  |  |
| 18 | TCONS_00016273 | OsGH3.1 | LOC_Os01g57610 | Enhance resistance | Ruan et al. 2019 |
|  | TCONS_00016274 |  |  |  |  |
|  | TCONS_00014126 |  |  |  |  |
|  | TCONS_00068710 |  |  |  |  |
|  | TCONS_00036122 |  |  |  |  |
| 19 | TCONS_00016273 | OsPGIP2 | LOC_Os05g01370 | Enhances resistance against pathogens by inhibiting fungal polygalacturonases | Kalunke et al. 2015 |
|  | TCONS_00016274 |  |  |  |  |
|  | TCONS_00016275 |  |  |  |  |
|  | TCONS_00014126 |  |  |  |  |
|  | TCONS_00068710 |  |  |  |  |
|  | TCONS_00036122 |  |  |  |  |

### 3.3 Characterizing the Expression Profile and Length Distribution of DE-lncRNAs across Pathogenic Infections

Figure 2 illustrates the expression profiles of DE-lncRNAs and their length and chromosomal distribution under various pathogenic infections. Figure 2A displays the length distribution of DE-lncRNAs in R-B, R-F, and R-V. The majority of DE-lncRNAs in R-B and R-F fall within the 200-1000 nt range, with varying percentages being upregulated and downregulated. However, same pattern is observed in uninfected rice samples (data not shown) suggesting that this length distribution profile is characteristic of lncRNA even under normal condition. In contrast in the case of R-V, a distinct pattern emerges, with a peak in longer DE-lncRNAs (1000-2000 nt and 2000-3000 nt ranges). Figure 2B shows the fold changes in expression levels of DE-lncRNAs, revealing consistent trends within the 200-7000 nt range for upregulated DE-lncRNAs. Notably, R-F exhibits ∼15 fold downregulation of DE-lncRNAs in the 200-1000 nt range, which is a significantly higher downregulation compared to R-B and R-V. Figure 2C illustrates the chromosomal distribution analysis, showing how the DE-lncRNAs are distributed across all chromosomes. Number of lncRNA’s mapped to each chromosome were further normalized to the chromosome size (in Mb) to obtain density distributions (Supplementary Figure 1). Upon normalization it is observed that there are variations in the DE-lncRNA densities across chromosomes in the different infection scenarios. In R-F, it is observed that Chromosome 8 has twice the density of lncRNA’s compared to chromosome 10. While, in R-V, notably it is observed that, Chromosome 12 is fourfold depleted and Chromosome 8 is twofold depleted in DE-lncRNA density compared to chromosome 10. In contrast in R-B, there is no significant difference in lncRNA distribution across the chromosomes with a +/- 20% variation. The functional origin and significance of these variations in DE-lncRNA density across the chromosomes is currently unknown and unexplained and could be explored experimentally in the future.

**Figure 2.**
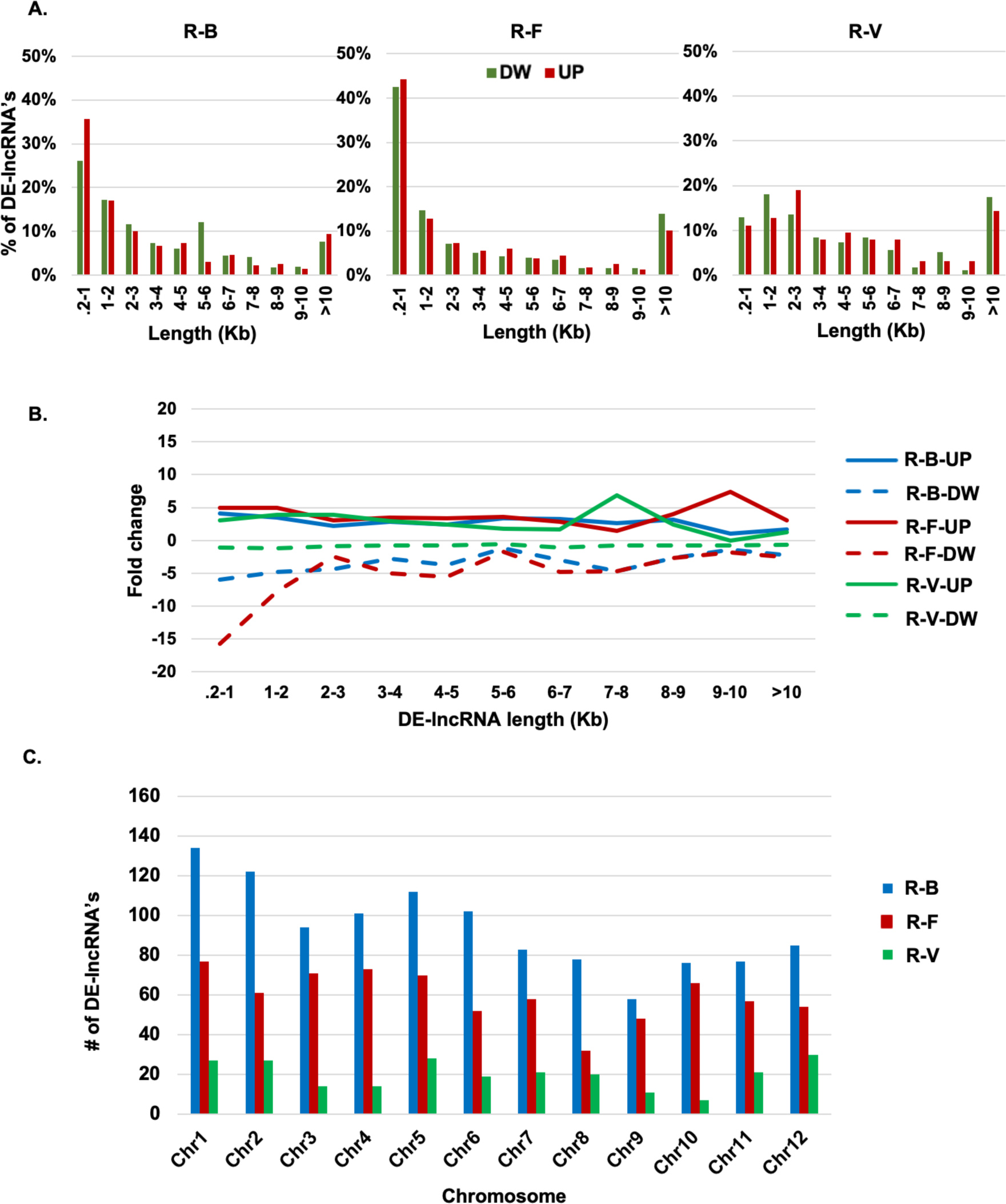
Differential expression profile of Rice lncRNA’s upon infection: **A.** DE-lncRNA length vs % occurrence, DW – Down Regulated. UP – Up Regulated; **B.** DE-lncRNA length vs Fold change in expression. **C.** Chromosomal mapping of differentially regulated lncRNA’s. R-B: Xanthomonas oryzae + CT9737-6-1-3P-M, R-F: Magnaporthe oryzae + LTH; and R-V: Rice black streaked dwarf virus + Wuyujing No. 7.

### 3.4 Chromosomal mapping of lncRNA and QTL

Using QTL analysis and chromosomal mapping, we identified potential genetic links between DE-lncRNAs and specific pathogenic infections. Figure 3 displays QTLs overlapping with DE-lncRNAs in R-B, R-F, and R-V infections, situated on Chromosomes 1, 3, 4, 5, 11, and 12. Notably, we found one common QTL co-localizing with a specific lncRNA across all three infections. Additionally, we detected six shared QTLs associated with 16 DE-lncRNAs in R-B and R-F infections, two QTLs correlated with three DE-lncRNAs in R-B and R-V infections, and two QTLs overlapping with four DE-lncRNAs in R-F and R-V infections. Interestingly, unique QTLs emerged for each infection type, as detailed in Supplementary Table 1. The shared QTLs have previously been linked to defence mechanisms, reinforcing the defence-related roles of many DE-lncRNAs identified in this study.

**Figure 3.**
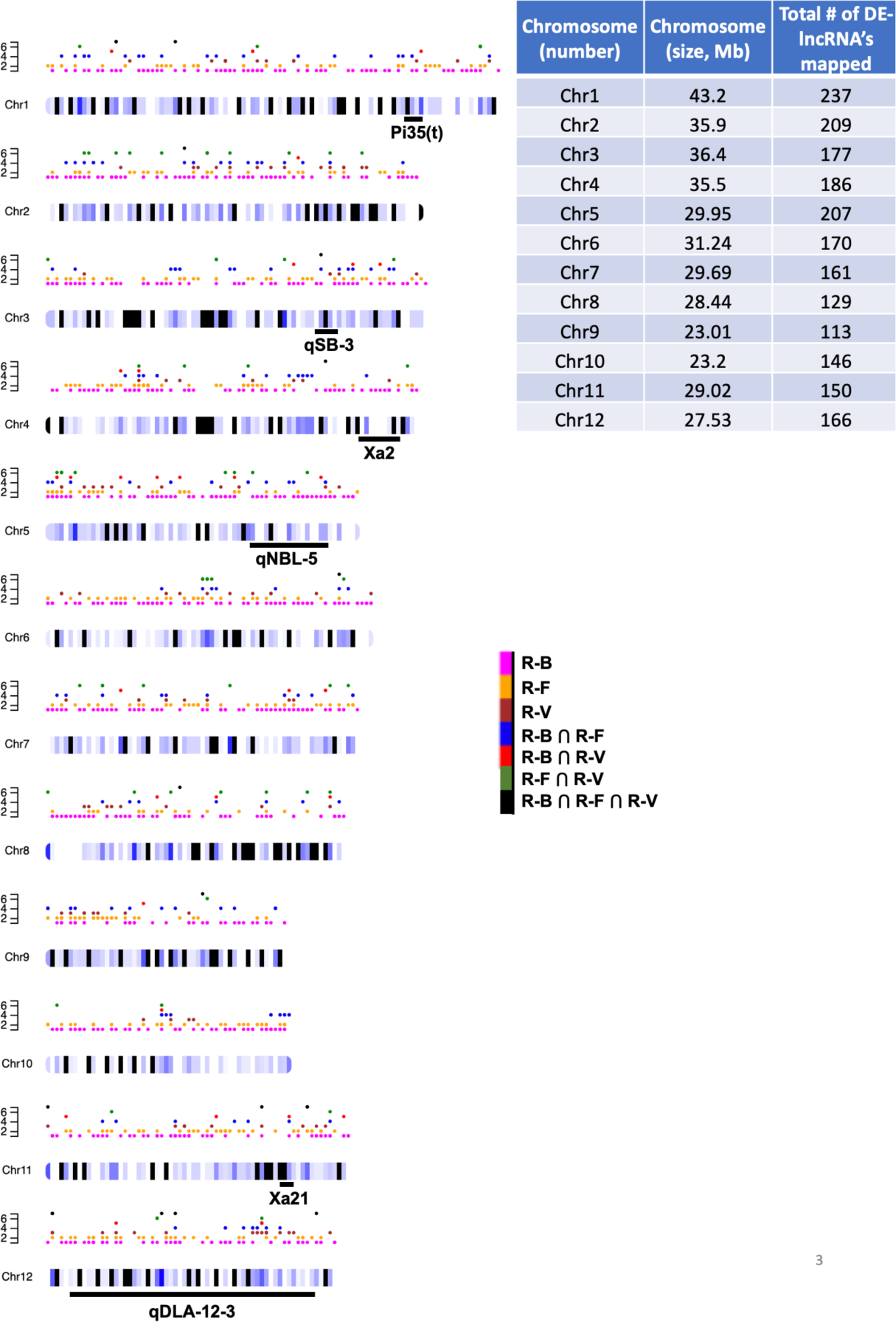
Chromosomal map of lncRNA and QTL: Each dot indicates different lncRNA, while the black horizontal lines below the chromosomes shows the QTL position. Different shades of blue on chromosome indicate the density of lncRNA, darker shade of blue indicate high density in comparison to lighter shade of blue while black indicate no lncRNA. lncRNA’s that are unique, shared and common between the 3 different infection pairs are depicted in differently colored dots. The scale bar on the left of each chromosome depicts the number of lncRNA’s of each type. All the QTLs indicated in the figure are related to blast or blight resistance function. R-B: Xanthomonas oryzae + CT9737-6-1-3P-M, R-F: Magnaporthe oryzae + LTH and R-V: Rice black streaked dwarf virus + Wuyujing No. 7. The table depicts the number of lncRNA’s mapped to each chromosome and compared with respect to the chromosome size.

### 3.5 Expression value calculation and PCA analysis

The expression levels of each transcript were quantified as transcripts per million (TPM) using raw read counts. As depicted in Figure 4, PCA analysis was carried out on differentially expressed lncRNAs and genes expressed under control and infection conditions. PCA analysis unveiled distinct infection-specific expression patterns. Specifically, in the case of R-B and R-F, the two states, infected vs uninfected control, exhibit a significant separation, whereas in R-V, there is a high overlap observed between the two states.

**Figure 4.**
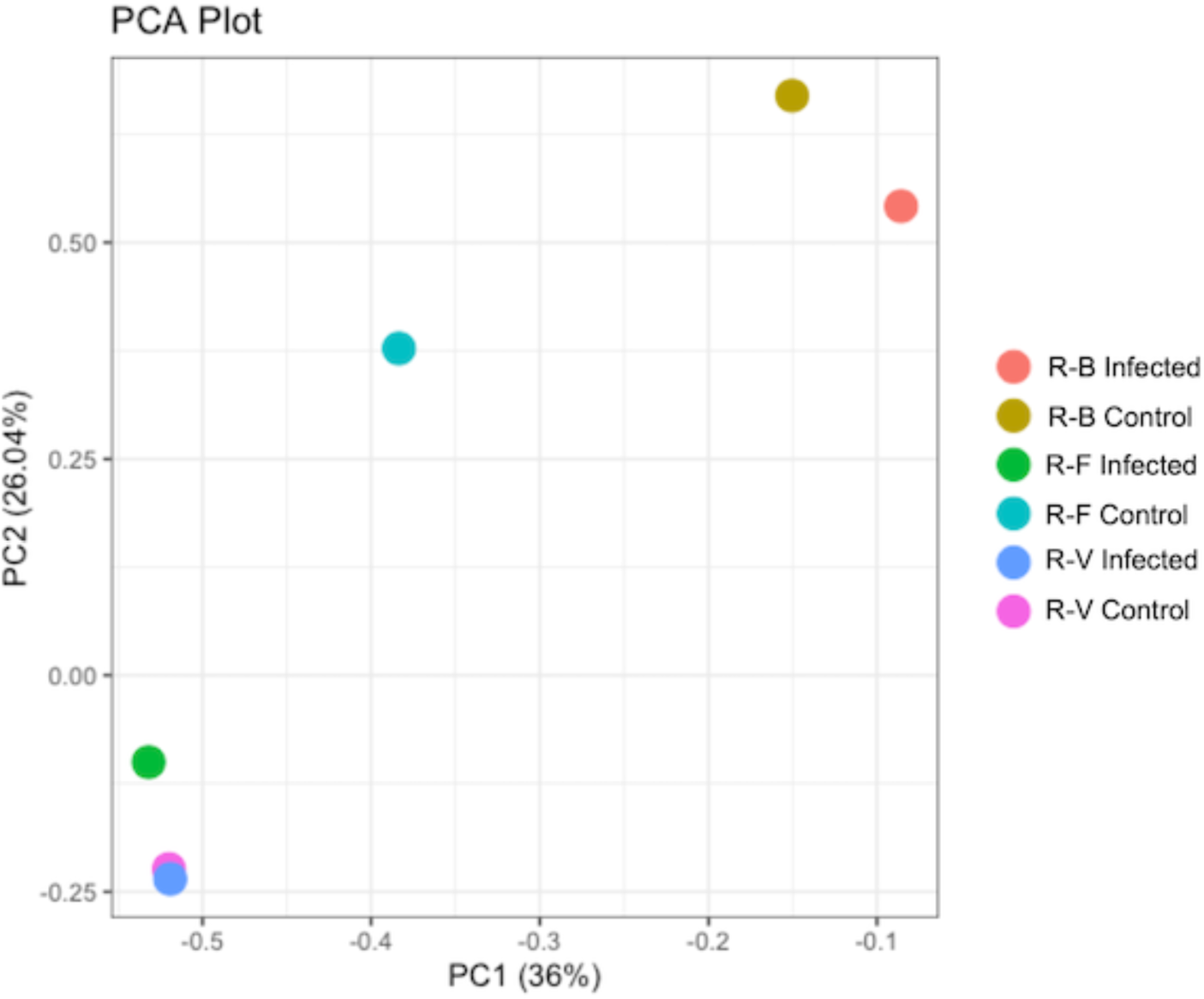
PCA Analysis of Control and Infected Samples: This figure presents the PCA analysis of control and infected samples for R-B, R-F, and R-V. The PCA plot visualizes the variance in the expression levels of differentially expressed genes (DE-genes) targeted by lncRNAs, allowing for the observation of potential clustering patterns. R-B: Xanthomonas oryzae + CT9737-6-1-3P-M, R-F: Magnaporthe oryzae + LTH and R-V: Rice black streaked dwarf virus + Wuyujing No. 7.

### 3.6 Prediction of Cis- and Trans-targets of DE-lncRNA’s: Revealing Common and Unique lncRNA Profiles and their Functional Analysis

LncRNAs manifest their effects via cis- or trans-regulation of their target DE-genes. Cis-regulation involves the regulation of protein-coding genes in proximal or overlapping regions, whereas trans-regulation suggests the regulation of DE-genes that are either located on the same chromosome at a further distance or other chromosomes. To determine their putative functions, we identified both cis and trans targets of the DE-lncRNAs.

#### 3.6.1 Prediction of DE-lncRNA target DE-genes and their functional analysis

In Figure 5A, we observe a total of 13,656 target DE-genes (6374 Cis-; and 7282 Trans-) targeted by 1124 DE-lncRNAs in R-B. In R-F, 691 DE-lncRNAs impact 3,532 DE-genes, with 2,457 undergoing cis-regulation and 1,075 exhibiting trans-regulation. Likewise, in R-V, 240 DE-lncRNAs target 1,519 DE-genes, comprising 334 cis-regulated and 1,185 trans-regulated DE-genes.

**Figure 5.**
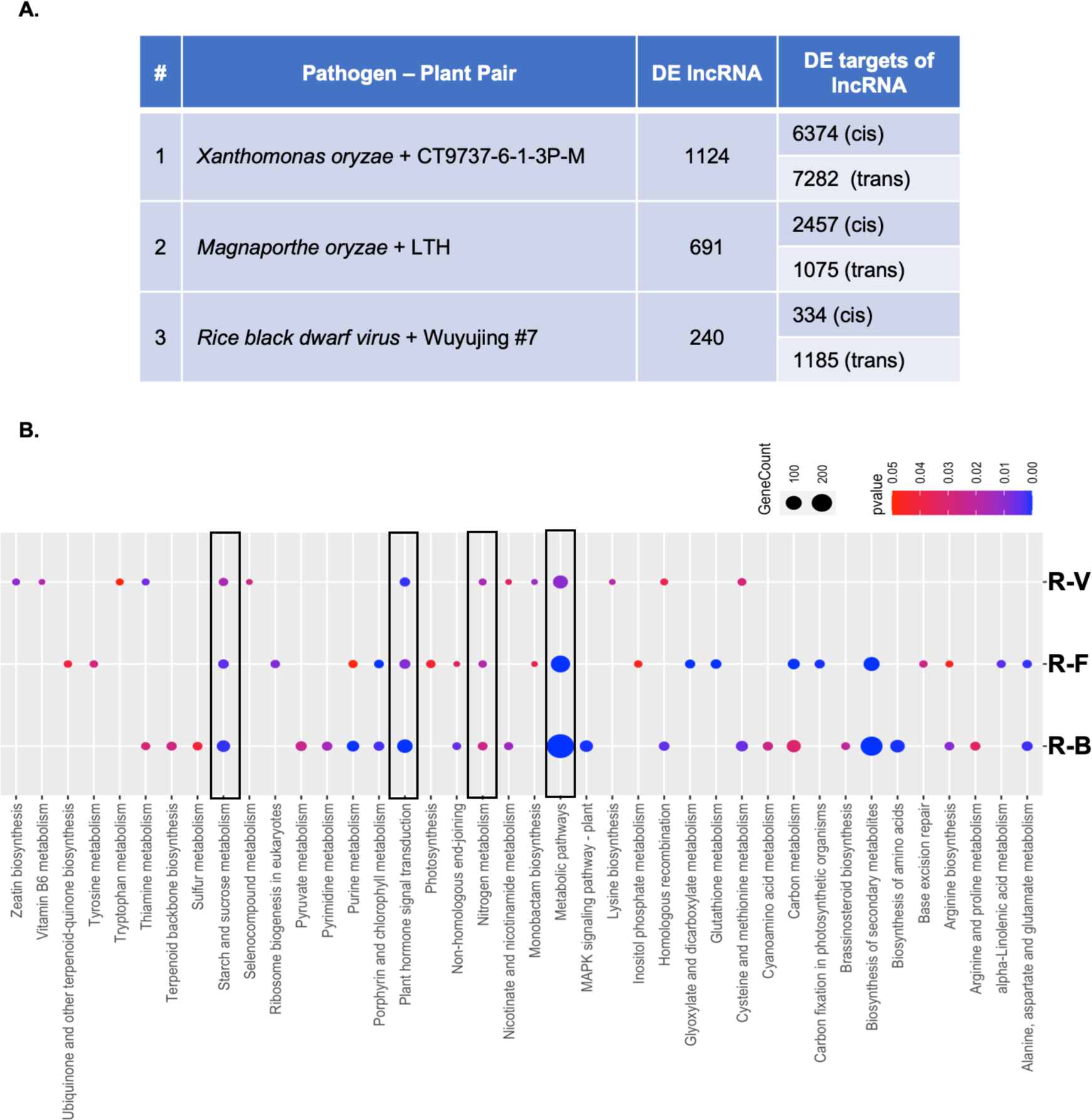
Total Identified lncRNAs and their Targets: **A.** Table lists the number of lncRNAs that are differentially expressed during bacterial, fungal, and viral infections, along with the number of target genes they regulate in cis and trans. **B.** This panel presents the results of KEGG pathway enrichment analysis for cis-trans target DE-genes of lncRNAs under different pathogen interactions (R-B, R-F, and R-V). The colour of the bubbles ranges from red to blue, with red indicating a significance level (P-value < 0.05) and blue indicating lower P-values and higher significance. The size of each bubble corresponds to the number of genes in each pathway. R-B: Xanthomonas oryzae + CT9737-6-1-3P-M, R-F: Magnaporthe oryzae + LTH and R-V: Rice black streaked dwarf virus + Wuyujing No. 7.

In Figure 5B, we conducted KEGG enrichment analyses for Cis- and Trans-target DE-genes in individual infections. R-B displayed 24 enriched KEGG pathways, R-F had 22, and R-V had 14. Four KEGG pathways (boxed in Figure 5B) — plant hormone signal transduction, metabolic pathways, starch & sucrose metabolism, and nitrogen metabolism—were common across all three infection scenarios. R-B and R-F shared seven KEGG pathways in common, including biosynthesis of secondary metabolites, purine metabolism, alanine aspartate and glutamate metabolism, porphyrin and chlorophyll metabolism, non-homologous end-joining, arginine biosynthesis, and carbon metabolism. R-B and R-V had four common KEGG pathways that include homologous recombination, cysteine and methionine metabolism, nicotinate and nicotinamide metabolism, and thiamine metabolism. Only one KEGG pathway, monobactam biosynthesis, was common between R-F and R-V. Additionally, unique KEGG pathways were identified for each infection, suggesting distinct plant responses to the infection mechanisms of the three pathogen types.

#### 3.6.2 Quantification of DE-lncRNAs and Analysis of Common and Unique DE-lncRNA-DE-gene Pairs

In Figure 6A, the Venn diagram illustrates unique and common DE-lncRNAs among three distinct plant pathogen pairs: R-B, R-F, and R-V infections. These three infections share 19 DE-lncRNAs in common. Furthermore, R-B and R-F have 155 lncRNAs in common, R-F and R-V share 36 lncRNAs, and R-B and R-V share 61 lncRNAs. In contrast, R-B, R-F, and R-V have 889, 481, and 124 unique DE-lncRNAs, respectively. Notably, among the 19 lncRNAs shared by all three infections, eight were downregulated, and one was upregulated in all infections. Additionally, six lncRNAs exhibited similar expression in both R-B and R-V, two in R-B and R-F, and two in R-F and R-V (Supplementary Table 2).

**Figure 6.**
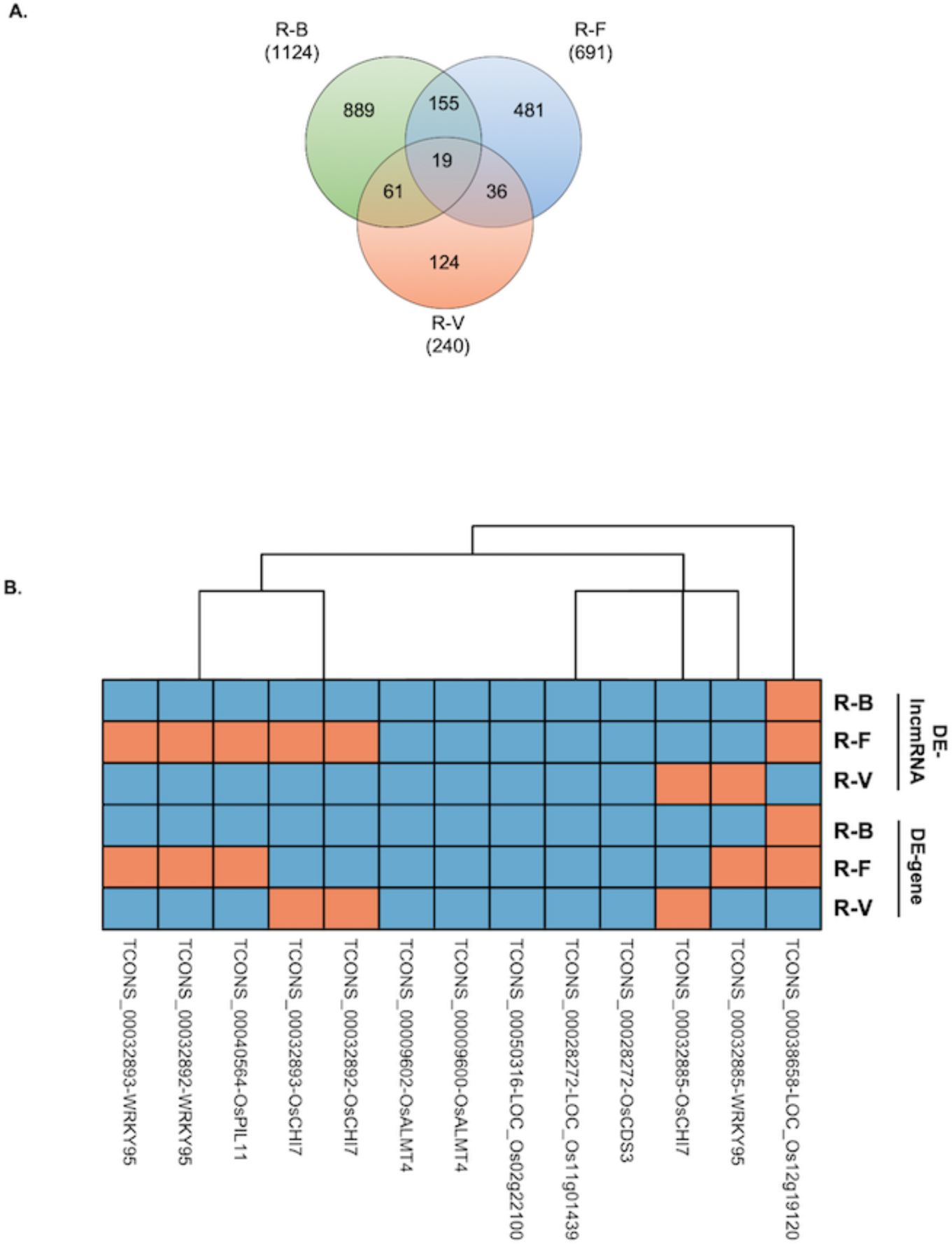
Analysis of differentially expressed lncRNAs and their target genes under different infection scenarios: **A.** Venn diagram depicting the common and unique DE-lncRNAs amongst the different infection scenarios **B.** Heatmap of the 13 common DE-lncRNA + DE-gene pairs under Bacterial, Fungal and Viral infection scenarios. Orange shows upregulation and green shows downregulation. R-B : Xanthomonas oryzae + CT9737-6-1-3P-M, R-F: Magnaporthe oryzae + LTH and R-V: Rice black streaked dwarf virus + Wuyujing No. 7.

We systematically analysed these 19 shared DE-lncRNA’s and their corresponding DE-gene pairs and identified that 13 of these ‘pairs’ are consistently modulated in response to all three infections (Figure 6B, Supplementary Figure 2). Of these, five exhibited uniform downregulation in all infections. Additionally, three pairs showed similar expression trends in R-B and R-V, while three displayed infection-specific patterns warranting further investigation. Furthermore, we found 511, 71, and 46 common pairs between R-B:R-F, R-B:R-V, and R-F:R-V, respectively (Supplementary Table 3, Supplementary Figure 2). Additionally, 29,796 pairs were exclusively differentially expressed in R-B, 6,302 in R-F, and 2,938 in R-V.

In summary, our in silico analysis identified 13 consistently altered DE-lncRNA-DE-gene pairs across all three infections, with some pairs exhibiting idiosyncratic trends. Additionally, a substantial fraction of DE-lncRNA-gene pairs were shared between two infection pairs, while some were uniquely expressed in each infection scenario. These findings offer valuable insights into the regulatory mechanisms of the studied plant-pathogen interactions. While our work is an in silico computational analysis, further exploration of infection-specific lncRNA-gene pairs can enhance our understanding of the functional roles of these lncRNAs in different pathogenic infections. This opens up avenues for experimental biology labs to explore such areas using techniques like RT-PCR, RNA-chip, and pull-down assays for further progress.

### 3.7 Cis- and Trans-target DE-genes of rice lncRNA’s are implicated in plant response to biotic stress

In Figure 7, we employed MapMan for the analysis of cis- and trans-regulated target DE-genes governed by DE-lncRNAs. Our analysis highlights the significant pathways influenced by DE-lncRNAs. These pathways include respiratory burst, signalling, hormonal pathways, and transcription factors, which in turn affect various plant defence-related processes. These pathways impact physiological processes such as cell wall formation, beta-glucanase activity, secondary metabolite production, and the expression of Pathogenesis-related (PR) genes.

**Figure 7.**
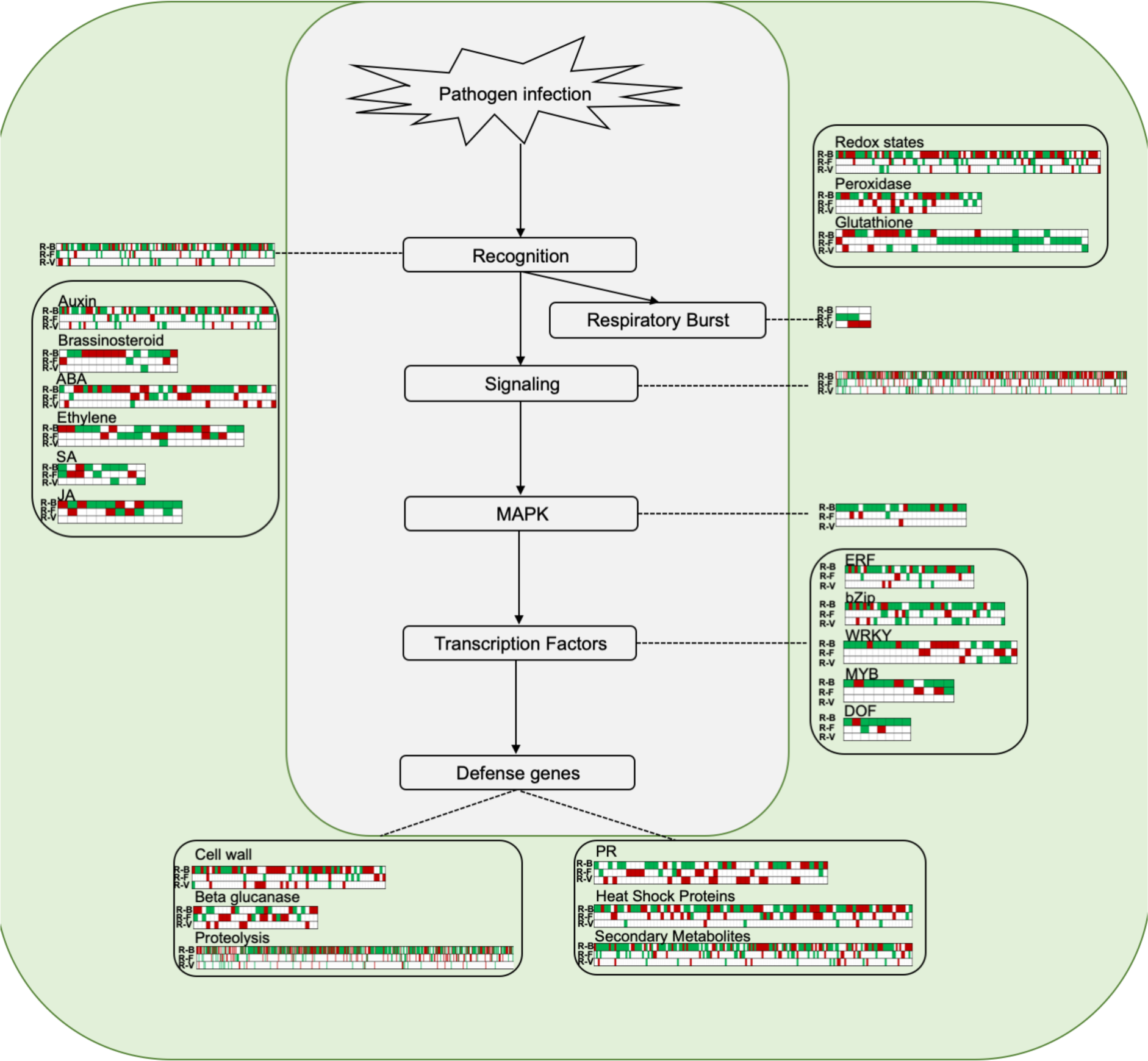
Overview of genes targeted by lncRNA under different pathogen infection scenarios: This figure illustrates the changes in expression of genes targeted by lncRNA in rice as a response to the biotic-stress of different pathogen infections: R-B (Xanthomonas oryzae + CT9737-6-1-3P-M), R-F (Magnaporthe oryzae + LTH), and R-V (Rice black streaked dwarf virus + Wuyujing No. 7). For each case, only genes with significant (P < 0.05) expression changes between infected and control samples, as identified by MapMan, are displayed. Upregulated gene expression is indicated in red, downregulated expression is shown in green.

## 4 Discussion

In recent years, there have been increasing evidence supporting the crucial role of lncRNAs in gene regulation during host-pathogen interactions. Studies have shown that lncRNAs are involved in the response of plants to various pathogens, including fungi, bacteria, and viruses (Huang et al. 2023).

Our in silico analysis identified DE-lncRNAs across these infections, employing the analytical pipeline detailed in Figure 1. The length distribution of DE-lncRNA’s (Figure 2A) observed in the case of R-B and R-F showed similar patterns, where the 200-1000nt length lncRNA’s showed maximum changes in their expression; whereas for R-V; the peak shifted to longer lengths in the 1000-3000nt range. This suggests a subtle difference in the nature of regulatory changes happening in the rice plant upon diverse infections.

We further analyzed whether there are any major differences between the different infection scenarios in the extent (amplitude of response) to which the DE-lncRNA’s are regulated. Notably, we observe that in the case of R-F, the DE-lncRNA’s of rice undergo a significant ∼15 fold downregulation in the 200-1000nt length range. This is not observed in the case of R-B / R-F. The mechanistic significance of this observation is under investigation. By knowing the individual identities of lncRNA’s in this 15-fold reduction scenario, it could be possible to understand why such a major downregulation is happening and its biological significance.

Our in silico investigation further examined the chromosomal distribution of DE-lncRNAs, revealing that they are dispersed across the rice genome rather than clustered / depleted in any specific chromosomal hotspots (Figure 2C). To substantiate the role of these identified lncRNAs in plant defence, we explored their overlap with QTLs known to confer resistance against pathogens in rice. The list of QTL’s was downloaded from Funrice gene database, and the overlap was checked using the chromosome wise coordinates of QTLs and DE-lncRNAs. The significant overlap observed highlights the role of these lncRNAs in enhancing plant resistance to diverse pathogenic infections (Figure 3). For instance, TCONS_00038658 which overlaps with qDLA-12-3 common across all infection is known for its role in Blast resistance. This qDLA-12-3 also contains other lncRNA’s that are common for R-B and R-F, R-B and R-V and R-F and R-V. This shows a crucial link between the role QTL qDLA-12-3 in various pathogenic infection apart from Blast.

The PCA analysis of the DE-lncRNAs and genes shows significant variation of transcript expression in rice responses to different pathogen infections (Figure 4). The findings clearly demonstrate that when compared to bacterial and fungal infections; virus-infected plants exhibit the least number of changes. To further dive deeper into the basis of these differences, we compared the DE-lncRNAs from different infections (Figure 5A) and found that only a few lncRNAs were common in all infections.

Using KEGG pathway enrichment analysis, we were able to identify four pathways that are common to all 3-infection scenarios (Figure 5B), namely metabolic pathways, nitrogen metabolism, plant hormone signal transduction, and starch & sucrose metabolism, all of which play important roles in plant-pathogen interactions.

Metabolic pathways are involved in the production and degradation of metabolites, which are essential for plant defence responses against pathogens (Kaur et al. 2022). Interestingly, pathogens also make use of these metabolic pathways to survive within the host.

When plants encounter biotic stress, they respond by reallocating their nitrogen reserves to produce signalling molecules like polyamines, GABA, and glycine betaine. This leads to a moderate rise in ammonia toxicity within plant cells. Consequently, the precise control of nitrogen metabolism proves indispensable in managing biotic stress, as it impacts virtually all physiological plant processes. Crucial role of nitrogen metabolism is highlighted by Yang et al. (2022) in their exploration of the intricate regulation of tobacco resistance through nitrogen metabolic rate variations.

Plant hormone signal transduction pathways are important for regulating plant growth and development, as well as for mediating plant responses to biotic and abiotic stresses (Di et al. 2017). In the context of plant-pathogen interactions, plant hormones such as salicylic acid, jasmonic acid, and ethylene are known to play key roles in modulating defence responses (Helliwell et al. 2016; Di et al. 2017; Ma et al. 2022).

Finally, starch and sucrose metabolism pathways are involved in providing energy and carbon skeletons for various cellular processes, including defence responses against pathogens (Tauzin and Giardina 2014). Sucrose, for example, has been shown to be involved in the regulation of plant immunity, while starch metabolism has been linked to the regulation of plant growth and stress responses (Tauzin and Giardina 2014).

Several pathways were found to be shared between R-B and R-F, R-B and R-V, and R-F and R-V, with a considerable number of pathways unique to each infection scenario (Figure 5B). This observation highlights the existence of distinctive plant responses to the varying mechanisms employed by the three diverse pathogen types.

In summary, the identification of these shared pathways through KEGG pathway enrichment analysis offers valuable insights into the intricate molecular orchestration governing plant-pathogen interactions. This knowledge concerning the conserved pathways affected during pathogen interactions can be further validated and potentially harnessed to innovate novel strategies for the development of pathogen-resistant crop varieties.

### 4.1 Deciphering the role of plant defence related DE-lncRNA common across all three infections

Upon further deciphering the DE-lncRNA + Gene pairs that were differentially expressed, we found that 13 pairs (Supplementary Figure 2) were common to all the different infection scenarios. Of these target DE-genes, many have been implicated for their role in plant defence (Figure 6B).

Investigating TCON_00032885 / 893 / 892 OsCHI7, reveals a complex regulatory pattern in different infections. Specifically, OsCHI7 displayed upregulation in the case of R-V but downregulation during R-B and R-F infections. In parallel, TCON_00032892/3 were upregulated in response to R-F and downregulation in R-V. Notably, TCON_00032885 was downregulated in both R-B and R-F infections, but was upregulated in the R-V, mirroring OsCHI7, emphasizing its direct regulatory role. This regulatory network is significant in understanding OsCHI7, that codes for chalcone isomerase (CHI) enzyme, crucial in flavonoid biosynthesis. In bacterial and fungal infections, flavonoids rapidly transport to the infection site, initiating a hypersensitivity reaction, limiting pathogen propagation (Blount et al. 1992; Dai et al. 1996; Beckman 2000). In the context of viral infections, flavonoids enhance plant resistance, as seen in their response to the cucumber green mottle mosaic virus in watermelon. This response is initiated by the induction of flavonoid biosynthesis through salicylic acid (SA), highlighting the versatile role of flavonoids in defending plants against various pathogens (Liu et al. 2023).

Another pivotal gene, OsPIL11 (phytochrome-interacting factor) regulating grain size, was targeted by upregulated lncRNA, TCON_00040564, in R-F, but downregulated in R-B and R-V. OsPIL11 interacts directly with the transcription factor OsSPL14 (SQUAMOSA promoter binding protein-like, SPL) with dual role in plant immunity and yield enhancement. It has been experimentally verified that mutations targeting the miRNA-binding site suppressing OsSPL14 expression, led to increase in crop yield, even in Mor infection (Wang et al. 2018). Mechanistically, OsSPL14 influences the WRKY45 promoter, enhancing plant resistance.

OsALMT4 (Aluminium-activated Malate Transporters 4) is downregulated in all three infections, mirroring its targeting lncRNA. Overexpression of this malate-permeable ion channel, relates to nutritional disorders. The ALMT protein family member AtALMT1 is upregulated in Arabidopsis in response to pathogen infection (Liepman and Olsen 2001; Luo et al. 2021), indicating a potential role for OsALMT4 in pathogen interactions.

Apart from common DE-lncRNA-DE-gene pairs in all infection we also deciphered the defence related DE-lncRNA-DE-gene pairs that are common across two infections: 1) Common between R-B and R-F; 2) Common between R-F and R-V; and 3) Common between R-V and R-B.

### 4.2 Exploring Shared DE-lncRNA-DE-Gene Interactions in R-B and R-F infections

A total of 511 pairs of DE-lncRNAs and DE-gene were found to be common between R-B and R-F (Supplementary Figure 2, Supplementary Table 3). Considering that both Xoo and Mor are hemi-biotrophic pathogens, this substantial number of shared lncRNA-gene pairs in rice infected by Mor and Xoo is along expected lines. Notably, among these pairs, three differentially expressed (DE) target genes are associated with the lncRNAs OsSWEET14, OsbHLH65, and OsSRO1a. These genes are known for their roles in bacterial leaf blight (Duy et al. 2021), rice blast (Shin et al. 2014), and bacterial blight (Kashihara et al. 2020), respectively.

OsSWEET14, a sugar transporter, linked to heightened Xoo susceptibility, exhibits increased expression leading to reduced leaf sugar content, enhancing Xoo resistance (Luo et al. 2021). Proper regulation of OsSWEET14 is vital for maintaining balanced sugar levels within the plant, thereby affecting its susceptibility to different pathogens. In our study, we noted that OsSWEET14 was upregulated in response to R-B infection but downregulated during R-F infection, and the lncRNA TCONS_00032659 targeting the OsSWEET14 gene was upregulated in both. This suggests a specific gene regulation pattern in response to pathogens, making it a compelling candidate for deeper exploration.

Another gene identified in the study is OsbHLH65, a basic Helix–Loop–Helix transcription factor that is phosphorylated by OsMPK3 (Shin et al. 2014), to induce different defence signal transduction pathways; indicating its role in plant pathogen interactions. This gene was observed to be upregulated in R-B and downregulated in R-F, but the lncRNA targeting it was upregulated in both infections.

OsAMTR1, a pyridoxal-5’-phosphate enzymes known as aminotransferases, is crucial in biotic stress (Kothari et al. 2016). This gene was upregulated in both R-B and R-F, with pathogen-specific expression exhibited by lncRNA with downregulation in R-B and upregulation in R-B. Overexpression of similar genes such as glyoxylate aminotransferase (AGT) and Glutamate : Glyoxylate Amino Transferase (GGAT) genes has been shown to enhance resistance to many pathogens (Liepman and Olsen 2001), indicating the potential positive effect of OsAMTR1 on plant defence.

Our study also identified lncRNAs targeting DE-genes in hormone signalling, including OsSRO1a (similar to RCD ONE1a), downregulated in R-B and upregulated in R-F. This gene suppresses JA signalling by negatively regulating OsMYC2 (Kashihara et al. 2020). Upregulation of JA signalling enhances plant resistance to hemibiotrophic pathogens, inducing resistance against rice bacterial blight (Kashihara et al. 2020).

### 4.3 Exploring Shared lncRNA-Gene Interactions in R-B and R-V Infections

In the context of shared gene pairs between R-B and R-V infections (Supplementary Figure 2), the study observed an intriguing pattern related to OsRR6, which is recognized as a negative regulator of cytokinin (CK) signalling (Hirose et al. 2007). OsRR6 exhibited an upregulation in R-B but downregulation in R-V infection, and the corresponding targeted lncRNA displayed a similar expression pattern. Despite limited information regarding OsRR6’s role in plant-pathogen interactions in rice, existing evidence suggests that increased cytokinin levels enhance pathogen resistance. Co-treatment with CK and salicylic acid (SA) in rice leaf blades upregulated pathogenesis-related genes, OsPR1b and PBZ1, which are dependent upon key regulators OsNPR1 and WRKY45 in the SA pathway (Jiang et al. 2013). Hence, investigating the impact of OsRR6-mediated cytokinin regulation on plant resistance to biotic stresses holds substantial scientific interest.

### 4.4 Exploring Shared lncRNA-Gene Interactions in R-F and R-V Infections

We explored common gene pairs in R-F and R-V(Supplementary Figure 2), among common pairs lncRNA targeting OsNCX1, belongs to the Sodium/Calcium exchanger (NCX) gene family in rice, regulating calcium ion (Ca2+) levels in various cell types. OsNCX1 contains both an NCX domain and an EF-hand domain, and it exhibits high expression levels across various developmental stages and plant parts (Singh et al. 2015). OsNCX1 is vital for diverse physiological processes (Giladi et al. 2016). This gene was downregulated in R-F and upregulated in R-V. The DE-lncRNA targeting were upregulated in R-F, and downregulated in R-V. These findings reveal distinct regulation of NADPH oxidase and sodium/calcium exchanger genes in response to different pathogens, suggesting their roles in complex signalling during plant-pathogen interactions.

### 4.5 Biotic stress related gene expression in plants during pathogen infection

Our investigation has revealed that lncRNAs exert control over several key genes implicated in the response to biotic stress. This connection was established by mapping the data from the Mapman database to identify cis- and trans-target DE-genes of lncRNAs. These encompass genes associated with pathogen recognition, those governing respiratory processes, signalling molecules, transcription factors, and genes central to defence mechanisms (Figure 7).

#### 4.5.1 Insights into Innate Immunity

Pathogen recognition genes of plants play a key role in detecting pathogen-associated molecular patterns (PAMPs), initiating innate immunity against various pathogens. In Arabidopsis thaliana, the NADPH oxidase gene, OsrbohD, plays a key role in recognizing PAMPs and generating reactive oxygen species (ROS) (Kadota et al. 2015). Our study revealed upregulation of OsrbohD in response to R-V infection and downregulation during R-F infection. Interestingly, the lncRNA targeting this gene showed the opposite pattern: downregulation in R-V and upregulation in R-F. These findings suggest that lncRNA-mediated regulation is critical in influencing ROS production during plant-pathogen interactions.

One intriguing and vital component of the plant’s innate immune response is the role of Transcription Activator-Like (TAL)-mediated immunity. This form of immunity involves the recognition of pathogen-specific effectors but operates independently of prior exposure to the pathogen. TAL effectors are unique to certain pathogens, yet plants recognize them as foreign invaders and mount a robust defence response when exposed to these effectors. For instance, the recessive form of the Xa5 gene has been identified as conferring resistance to bacterial blight (Iyer-Pascuzzi et al. 2008a, b). Xa5 is a TFIIA transcription initiation factor, and it is triggered by TAL transcription factors. Intriguingly, in our study, we observed that both the Xa5 gene and the DE-lncRNA targeting it exhibited differential regulation in response to different infections. Specifically, they were downregulated in the case of R-B and upregulated in response to R-F. Xoo TAL effectors play a dual role in either contributing to disease progression or triggering the plant’s defence mechanisms by binding to host DNA and activating effector-specific host genes (Chen and Ronald 2011). These findings suggest that the plant’s response to bacterial and fungal infections may involve distinct regulatory mechanisms and underscore the intricate and multifaceted nature of plant-pathogen interactions, which continue to be a subject of in-depth investigation in the field of plant biology.

RC24, a class I chitinase, hydrolyses fungal cell wall chitin, contributing to defence against fungal infections (Xu et al. 1996). RC24 was upregulated in R-F and R-V infections. In R-F, DE-lncRNAs targeting RC24 were upregulated, indicating coordinated regulation. In R-V, RC24-targeting lncRNAs were downregulated, suggesting distinct regulatory patterns during viral infections.

#### 4.5.2 ROS Regulation and Immune Responses in Plant-Pathogen

Reactive oxygen species (ROS) are pivotal in activating immunity against pathogens, especially under unfavourable conditions like biotic (Sahu et al. 2022) and abiotic (Gill and Tuteja 2010) stress, waterlogging (Guan et al. 2019), and nutrient imbalances (Görlach et al. 2015). ROS regulates biological processes, such as pathogen defence, programmed cell death, and stomatal control (Gill and Tuteja 2010). They collaborate with epigenetic modifiers and phytohormones, influencing plant development and stress responses (Tsukagoshi et al. 2010; Gill and Tuteja 2010; Zhang et al. 2017; Kong et al. 2018).

We categorized genes into three groups: Redox pathway genes (e.g., Thioredoxins and Ascorbate-Glutathione system), Peroxidases, and Glutathione S transferases.

Thioredoxins (Trxs) play a vital role in maintaining redox balance through thiol-disulfide exchange reactions, which activate defence-related proteins, including transcription factors (Zhou et al. 2023) and activation of defence pathways in response to pathogen infections. One such essential gene, OsTrxm, is a thioredoxin involved in the regulation of ROS levels. In the case of R-F infection, OsTrxm was downregulated, while the lncRNA targeting this gene was upregulated, suggesting potential lncRNA-mediated regulation of OsTrxm, which is crucial for an effective immune response (Yong et al. 2008).

Conversely, another gene, OsPrx30, classified as a Class III peroxidase, showed increased expression in response to R-F, along with its regulatory lncRNA. OsPrx30 acts as a scavenger of ROS (Liu et al. 2021) and its upregulation renders the plant more susceptible to pathogen infections. Similarly, OsGST4, a glutathione S-transferase protein known for its ROS-scavenging activity (Xu et al. 2018), exhibited upregulation in response to R-F. The study identified other additional genes within these categories, although their specific roles in plant-pathogen interactions remain uncharacterized. This opens new avenues for exploring novel genes and lncRNAs that may act as regulatory partners, working together to regulate plant immunity effectively.

### 4.6 Pathogen Recognition and Immune Signalling Pathways

Upon sensing a pathogen, plants activate several downstream signalling pathways. For example, CRK6 a cysteine-rich-receptor-like kinases gets upregulated on detection of pathogen, activating NH1-mediated immunity, enhancing pathogen resistance (Chern et al. 2016). This gene was upregulated in case of R-B, which shows its role in bacterial recognition. Furthermore, our analysis has revealed that CRK6, is targeted by three DE-lncRNA, TCONS_00097445, TCONS_00023604 and TCONS_00104360. Among these, TCONS_00097445 and TCONS_00023604 exert a trans-regulatory influence on CRK6, while TCONS_00104360 functions in a cis-regulatory manner. It is intriguing to note that all three of these DE-lncRNAs also display upregulation specifically in response to R-B infection, suggesting involvement of these DE-lncRNA’s in the plant’s defence mechanisms.

The mitogen-activated protein kinase (MAPK) pathway is central in plant defence. OsMAPK3, OsMAPK6, OsMPK7, and OsMAPK15 play roles in defence responses (Song and Goodman 2002; Li et al. 2019; Hong et al. 2019; Zhou et al. 2019). In our study, it was observed that expression of OsMAPK3, OsMAPK6, and OsMPK7 was downregulated, whereas OsMAPK15 was upregulated in response to the experimental conditions associated with R-B. Limited DE-lncRNA-regulated gene expression was observed in R-F and R-V, implying distinct regulatory mechanisms in response to different stressors. This complexity highlights distinct plant’s response to different stressors.

### 4.7 Navigating Plant Hormones: Exploring the Impact of lncRNAs on Defence

Plants use phytohormones like Salicylic Acid (SA), Jasmonic Acid (JA), Indole-3-Acetic Acid (IAA), and Ethylene (ET) for signalling in defence (Figure 7). These hormones play vital roles in orchestrating rice plants’ responses to pathogens.

SA triggers defences by inducing defence-related genes, leading to pathogenesis-related (PR) protein synthesis. Recent study by Wang et al. (2022) shows SA’s upregulation during infection by the fungal pathogen *Magnaporthe oryzae*. In the same way, OsBISAMT1 which encodes for S-adenosyl-L: -methionine:salicylic acid carboxyl methyltransferase was induced in case of blast fungus, *Magnaporthe grsiea* infection (Xu et al. 2006). In this study, OsBISAMT1 was downregulated in both R-B and R-V and the corresponding targeting DE-lncRNAs were also downregulated, no DE-lncRNA regulation was observed in R-F. This research highlights the potential involvement of lncRNAs in fine-tuning the SA signalling pathway, thereby warranting further in-depth study.

JA regulates defences by activating genes for antimicrobial compounds and protease inhibitors. OsJAR1 makes major contributions to stress-induced JA-Ile production and is involved in defence responses in rice leaves (SHIMIZU et al. 2013). In this study, OsJAR1 was downregulated in R-B, so is the DE-lncRNA targeting it. These findings are in agreement with the putative role of TCONS_00090193 in modulating JA-based defence mechanisms in rice.

IAA has a multifaceted role in plant immunity, affecting growth and development. The auxin responsive genes of GH3 family regulates growth and development and, hence, are deeply involved in a broad range of physiological processes. Overexpression of OsGH3.1 in rice enhances resistance to *Magnaporthe grisea* infection (Domingo et al. 2009). OsGH3.1 was downregulated in R-F in our study, along with its targeting lncRNA.

ET contributes to rice’s defence against the bacterial pathogen *Xanthomonas oryzae* pv. oryzae (Xoo). ET triggers defence responses, including antimicrobial compound synthesis and cell wall reinforcement (Zhang et al. 2016). In our study we identified number of DE-lncRNA targeted differentially expressed genes related to ET, but none of them have been experimentally characterized for their role in plant pathogen interaction, thus giving a enormous scope for experimentally characterizing their function.

In summary, SA, JA, IAA, and ET govern the intricate relationship between rice plants and pathogens, influenced by the pathogen type and signalling pathways involved. Understanding these interactions enhances understanding of plant immunity and offers strategies for disease management in rice.

### 4.8 Modulation of Transcription Factor Responses to Pathogenic Microorganisms and Activation of Protective Genes

Transcription factors (TFs) are integral to plant defence mechanisms and play a crucial role in orchestrating plant responses by mediating a molecular dialogue between the plants and pathogens. TFs mediate effector-triggered immunity (ETI) and PAMP-triggered immunity (PTI) (Zhang and Zhou 2010), the two primary branches of plant immune responses.

In the current study, we analysed ERF, bZip, WRKY, MYB, and DOF transcription factor families (Figure 7), revealing distinct expression patterns in response to various pathogens. This highlighted their diverse roles in plant defence, potentially linked to specific pathogenic cues.

An intriguing aspect is the emerging connection between lncRNA’s and TFs. Although the exact nature of this interaction remains mysterious, preliminary findings suggest that lncRNA’s may modulate TF activity in response to pathogens. Interestingly, TF regulation by DE-lncRNA’s was most pronounced during R-B infection compared to R-F and R-V, opening exciting avenues for understanding plant immunity mechanisms.

Exploring the crosstalk between lncRNA’s and TFs in plant-pathogen interactions holds promise for future research.

### 4.9 Directing Plant Defences; Activation of Protective Genes

The aim of plant defence signalling pathways is to activate specific defence genes, which fall into two main groups. Group 1 includes genes related to cell wall, beta glucanase, and proteolysis. Group 2 comprises pathogenesis-related (PR) genes, heat shock proteins, and secondary metabolites (Figure 7).

Within Group 1, genes mainly impact the pathogen’s cell wall composition. For instance, OsPG1 codes for Polygalacturonase, influencing cell wall architecture, programmed cell death suppression, and cell wall integration, thereby regulating plant immunity (Cao et al. 2021). In our study, we made a noteworthy observation of the downregulation of OsPG1 in response to both R-B and R-V infections. This downregulation was accompanied by a corresponding decrease in the expression of DE-lncRNAs, with TCONS_00116715 in R-V and TCONS_00106037 in R-B exhibiting similar downregulated trends. Conversely, OsPGIP2, which encodes a polygalacturonase-inhibiting protein, enhances resistance against pathogens by inhibiting fungal polygalacturonases (Kalunke et al. 2015). Our investigation showed downregulation of OsPGIP2 in R-F, increasing susceptibility to Mor infection. Studying the regulatory mechanisms of OsPGIP2, particularly involving lncRNA’s, can provide deeper insights.

In Group 2, genes like OsOSM1 enhances rice resistance against sheath blight, are upregulated in response to jasmonic acid (JA) (Xue et al. 2016). In our study OsOSM1 was upregulated in R-B, and the DE-lncRNA targeting it was also upregulated. This observation strongly suggests that OsOSM1 may play a significant role in countering bacterial infections and that its regulation is influenced by DE-lncRNAs. This also highlights the importance of understanding the regulatory networks controlling Group 2 genes for enhanced disease resistance in plants.

## 5 Conclusion

This study aims to investigate the roles of lncRNAs in the interactions between plants and various pathogens, specifically bacterial (R-B), fungal (R-F), and viral (R-V) infections in rice. Our in silico analyses revealed distinct responses to these pathogenic challenges. Notably, viral infections induced fewer gene expression changes compared to bacterial and fungal infections, emphasizing the need for targeted strategies to protect rice crops. We also explored how lncRNA’s regulate key genes such as OsCHI7 and OsSWEET14, which play vital roles in defence mechanisms and pathogen recognition. Furthermore, we examined the involvement of transcription factors like CRK6 in immunity pathways and the importance of balanced reactive oxygen species (ROS) production in immune responses. Overall, our in silico study identified novel lncRNA that specifically interact with genes involved in pathogen recognition, signalling, and defence. These identified lncRNA and gene candidates are ripe for further experimental validation, which could provide valuable insights into their role in plant-pathogen interactions and their influence on defence and signalling pathways.

## Conflict of interest

The authors declare no conflict of interests.

## Acknowledgements

MK, MS and BR acknowledge Jio Platforms Limited, for providing the necessary facilities required and financial assistance through the salaries provided.

---

**Supplementary Figure 1.**
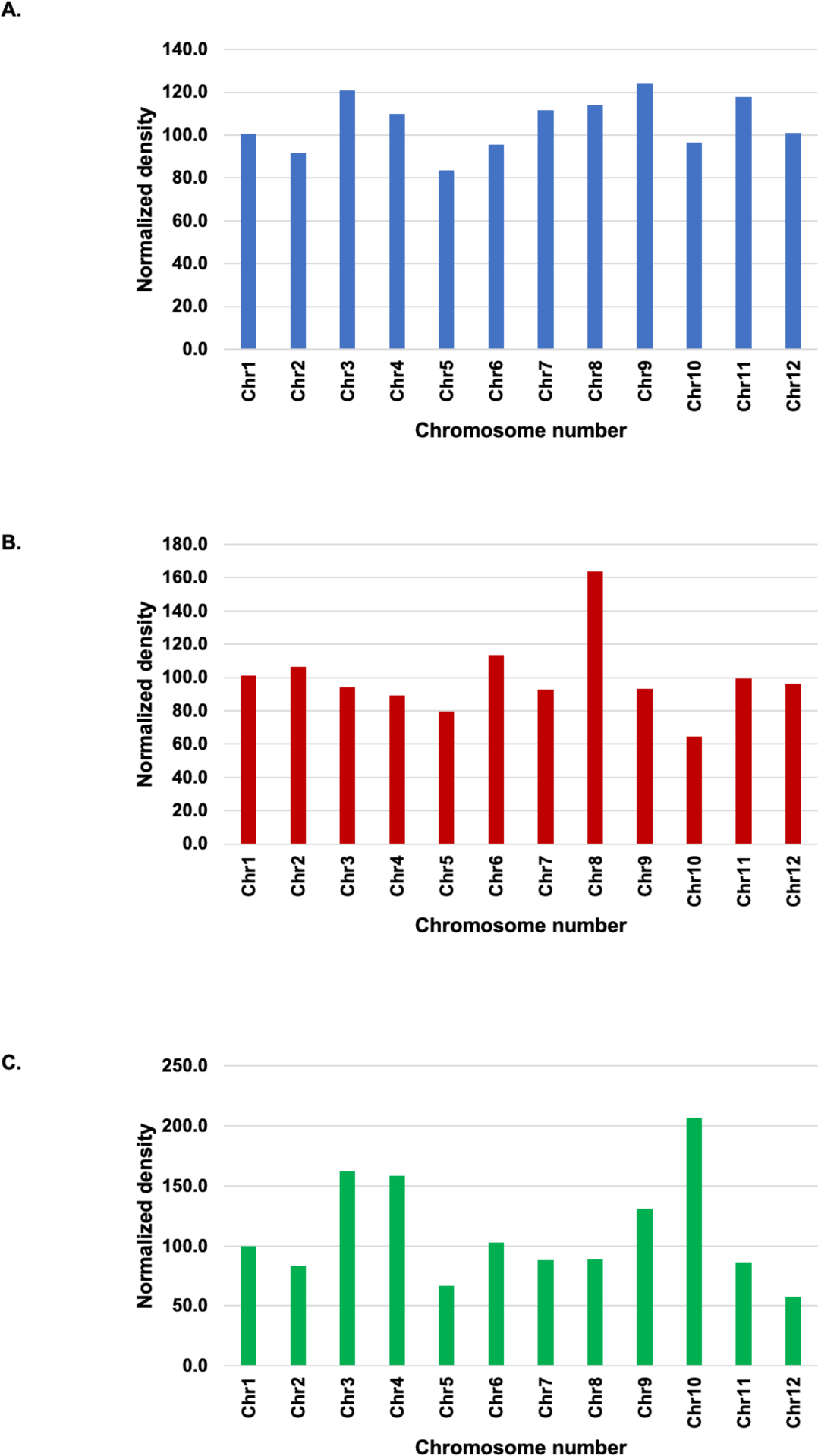
Distribution of DE-lncRNAs on chromosomes normalized to chromosome length. The figure illustrates the normalized distribution of DE-lncRNAs across chromosomes in response to distinct pathogenic infections. Initially, the lncRNA density was determined by dividing the length of each chromosome by the number of lncRNA present on the chromosome. Subsequently, the lncRNA densities were normalised using the chromosome 1 density as 100%. Each panel corresponds to a specific infection condition: **A)** R-B: Xanthomonas oryzae + CT9737-6-1-3P-M **B)** R-F: Magnaporthe oryzae + LTH; **C)** R-V: Rice black streaked dwarf virus + Wuyujing No. 7

**Supplementary Figure 2.**
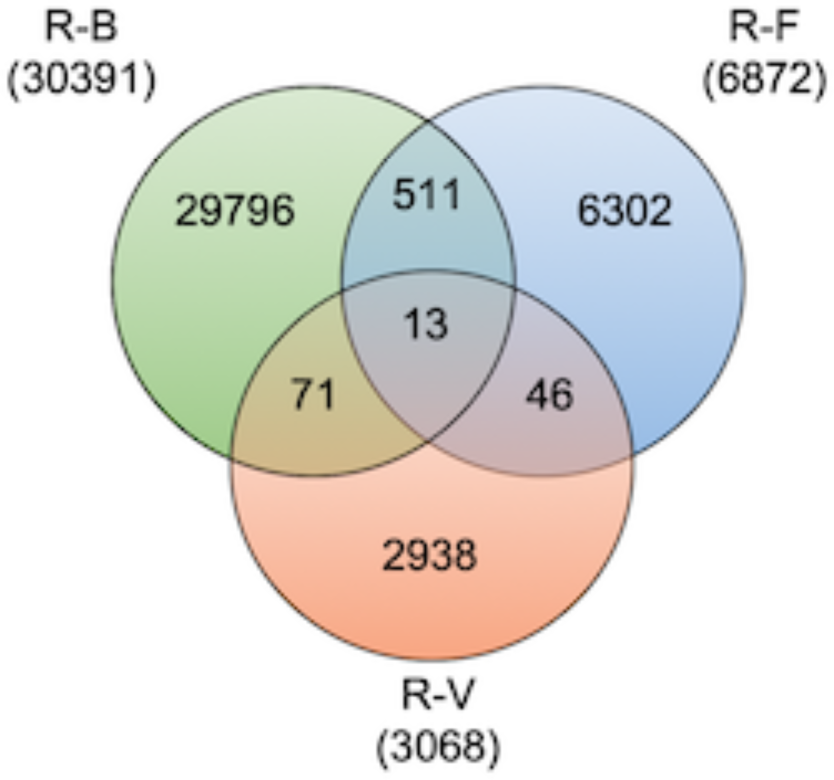
Venn diagram depicting the common and unique DE-lncRNAs-gene pairs amongst the different infection scenarios.

## Notes

### Competing Interest Statement

The authors have declared no competing interest.

